# Discovering Novel PETase Enzymes for Enhanced PET Plastic Degradation Using in Silico Approaches

**DOI:** 10.64898/2026.02.12.705489

**Authors:** Bhagyashree Patil, Arbaz Attar, Amrendra Kumar, Satyabharati Giri

## Abstract

Accounting for 12% of global solid waste, Poly (ethylene terephthalate) (PET) is one of the most abundantly produced synthetic polymers. While PET offers substantial commercial benefits, its widespread use has led to disproportionate environmental hazards due to its resistance to degradation. To address this problem, several solutions have been proposed, including enzymatic degradation via PETase, MHETase, and Cutinase. Among these, PETase exhibited significant PET-degrading activity. However, the application of PETase has been hampered by its lack of robustness to pH, temperature ranges, and slow reaction rates. Hence, it has become novel enzymes that can overcome these limitations and function efficiently. In this study, we utilised an integrated in silico bioinformatics pipeline to identify and characterise novel PETase candidates from the *Thermophilic actinobacteria Thermobifida cellulosilytica* and *Thermobifida halotolerans* species. The PlasticDB database contains 228 plastic-degrading enzyme sequences. In which PETase (00188) is significantly homologous with two putative proteins, Hydrolase (ALF00495.1) and hypothetical protein (WOZ56011.1). The discrete optimized protein energy (DOPE) scores, stereochemical assessments, and homology modeling results closely mirrored our findings for both proteins, supporting their structural stability. The molecular dynamics simulations revealed that the putative ALF00495 variant exhibited more extensive and robust hydrogen-bonding networks, enhanced conformational stability, and increased structural compactness compared to the reference enzyme. The present in silico investigation underscores the potential of putative ALF00495 as a highly effective PETase biocatalyst for polyethylene terephthalate (PET) degradation. Collectively, these findings illustrate the utility of computational approaches of novel PET-degrading enzymes, thereby facilitating the development of sustainable biotechnological strategies to mitigate global plastic pollution.

## Introduction

A century ago, the invention of the first fully synthetic plastic marked the beginning of a material revolution. What started as a scientific breakthrough soon transformed daily life, with plastics woven into nearly every aspect of modern society [1]. Their durability and affordability made them indispensable, from packaging our food to building our homes and supporting advances in healthcare and engineering[2]. Yet, the very qualities that made plastics so valuable have also become their greatest curse. Resistant to decay, they now accumulate as a major global pollutant, threatening ecosystems and driving environmental crises[3]. Nowadays, the world’s most common plastics—polyethylene, polypropylene, polyvinyl chloride (PVC), polystyrene, and polyethylene terephthalate (PET) continue to shape industries while simultaneously challenging us with their long-term ecological consequences[4, 5]. Among these plastics, PET is a thermoplastic polymer that contributes significantly to the global accumulation of plastic waste each year. Due to its chemical stability, hydrophobicity, and high molecular weight, PET is exceptionally resistant to biodegradation, persisting in landfills and aquatic environments for centuries [6]. Recent estimates indicate that more than 70 million tons of PET are produced worldwide annually, of which less than 30% is effectively recycled, resulting in its accumulation in natural ecosystems and contributing to micro plastic pollution, biodiversity loss, and long-term ecological damage [7, 8].

The society has relied on traditional recycling methods to manage plastic waste, but these approaches come with heavy trade-offs. Mechanical recycling, though widely used, slowly wears down the very material it seeks to preserve, producing lower-quality plastics with each cycle [9–11]. Chemical recycling promised a more complete solution, yet its dependence on harsh reagents and enormous energy demands has kept it far from being viable at scale. Faced with these shortcomings, nature offered an unexpected ally [12]. Microorganisms, armed with specialised enzymes, can break down PET into its basic building blocks, terephthalic acid (TPA) and ethylene glycol (EG) [8, 13–15]. Unlike conventional methods, this biological strategy not only dismantles plastic waste but also makes it possible to recover monomers for reuse in new polymer synthesis, opening the door to a more sustainable cycle of plastic production and degradation [16].

Among the many enzymes uncovered that can degrade plastics, PETases have drawn the most attention as nature’s own specialists, though only a handful have been explored in detail. The PETase from *Ideonella sakaiensis* is perhaps the most relied for its surprising ability to break down PET at moderate, mesophilic temperatures[17]. This discovery came with several limitations, including quickly faltering catalytic power and stability under the harsher conditions required for industrial use. Other enzymes, such as cutinase-like hydrolases from *Thermobifida fusca* and related actinomycetes, also showed promise, but like the *I. sakaiensis* PETase, they need significant engineering before becoming commercially viable[18, 19]. In this context, thermophilic microorganisms emerge as promising candidates to drive breakthroughs in PET degradation. Thermophilic microorganisms thrive in high-temperature environments, and their enzymes exhibit inherent resistance to denaturation, making them attractive candidates for industrial biocatalysis. Elevated temperatures not only maintain enzymatic stability but also enhance PET chain mobility, thereby improving substrate accessibility for hydrolysis [20]. Within this framework, the genus *Thermobifida*, long recognized for its role in biomass degradation, has recently gained attention as a potential reservoir of novel PET-degrading enzymes [21, 22]. While *T. fusca* has received attention, its close relatives—*T. cellulosilytica* and *T. halotolerans* remain virtually unexplored. The limited diversity of currently known PETases, together with an incomplete understanding of their activity, continues to pose a challenge to advancing enzymatic PET degradation toward scalable applications [23].

To address this gap, we employed comprehensive bioinformatics approaches to explore and characterize potential PET-degrading enzymes from these underexplored *Thermobifida* species, thereby advancing the understanding of their sequence–structure–function relationships. In this study, advanced bioinformatics tools and curated databases have opened new avenues for the discovery of plastic-degrading enzymes through in silico methods [24–26]. PlasticDB [27] (https://plasticdb.org/) consolidates experimentally verified and computationally predicted plastic-degrading enzymes, serving as a valuable resource for mining functional sequences. When combined with sequence alignment tools, domain prediction servers, and homology modeling techniques, researchers can now conduct comprehensive virtual screens of the environment. A curated 228 protein sequences were retrieved from PlasticDB, encompassing a diverse range of hydrolases, esterases, cutinases, and PETase-like proteins. These sequences were subjected to BLASTp [28, 29] analysis against the RefSeq protein database to identify homologous proteins with significant sequence similarity to known PETases (00188). Candidate sequences were filtered based on stringent criteria, including sequence identity, alignment length, and the presence of conserved catalytic motifs such as the Ser–His–Asp triad, which is essential for hydrolytic activity [30]. Among the identified sequences, proteins from *T. cellulosilytica* and *T. halotolerans* emerged as promising candidates, representing novel sources of potential PETase activity. Further, structural modeling employs known PETase and cutinase structures as templates to generate high-confidence three-dimensional models[31]. These models were evaluated using stereochemical quality checks and scoring metrics such as the DOPE (Discrete Optimized Protein Energy) score. Structural superimposition with reference PETases helped confirm the presence and spatial alignment of key catalytic residues, further supporting their potential function. Additionally, to assess their stability and dynamic behavior under physiological conditions, molecular dynamics (MD) simulations were conducted using GROMACS [32] and root-mean-square deviation (RMSD), root mean square fluctuation (RMSF), analysis of solvent accessible surface area (SASA), hydrogen bond, and radius of gyration were calculated. The conformational flexibility of the enzyme models over time indicates favorable structural characteristics. These methods not only accelerate the discovery pipeline but also reduce the cost and time associated with wet-lab screening to demonstrate the potential of *T. cellulosilytica* and *T. halotolerans* as previously not well-recognised sources of PET-degrading enzymes. By leveraging an integrated in silico pipeline to identify novel candidate PETases with favorable properties for biotechnological application. These enzymes may serve as starting points for protein engineering or direct use in PET bioconversion systems. Ultimately, this research contributes to the growing toolkit for enzymatic plastic degradation and supports global efforts toward sustainable waste remediation and circular bioeconomy.

## 2. Materials and Methods

### 2.1. Data Collection

A total of 228 sequences were downloaded in FASTA format from plasticDB (https://plasticdb.org/)[27] database, a curated resource compiling known and predicted plastic-degrading enzymes. These sequences represented a variety of enzyme classes, including hydrolases, esterases, cutinases, and PETases, which are known or predicted to be active against PET (00188) and other synthetic polymers. The sequences were manually curated and filtered to remove incomplete entries and redundancies, forming the primary dataset for subsequent analysis.

### 2.2. Protein sequences Identification

BLASTp analysis was performed through the Galaxy server (accessed on 24 June 2025), utilizing the RefSeq Protein database (version updated on 3 August 2023) to identify homologous sequences [33] corresponding to established plastic-degrading enzymes. Furthermore, Homology was considered significant with an identity threshold of ≥70%, alignment length ≥100 amino acids, and e-value ≤1e-5. Sequences were identified using criteria for downstream structural modeling, and cluster analysis helped identify unique sequences linked to PET-degrading proteins in *T. cellulosilytica* and *T. Halotolerans* [26]. Additionally, the conservation of catalytic motifs, particularly the Ser–His–Asp catalytic triad, was manually evaluated within the alignment outputs to infer potential hydrolytic activity [34].

### 2.3. Structure Prediction and Model Building

Selected following proteins sequences (AEW43689, AHY26884, AJA43734, ALF00495, AOZ64398, ASR86604, ASR86801, AYB69015, AYN57234, CAM32977, QBJ00759, QFP94845, WOZ56011, WOZ56057) were used to construct three-dimensional models using the Modeller software [35] (Version 10). Structural templates were selected based on sequence similarity and functional relevance, with a focus on cutinase and PETase structures from *Thermobifida fusca* (PlasticDB ID: 00057) and *Ideonella sakaiensis* (PlasticDB ID: 00188) [36]. These template alignments were optimized, and multiple models were generated for each candidate. The final models were selected based on energy scores and refined to correct local distortions in loop regions and side chains [37].

### 2.4. Structure Validation and Scoring

The quality of the predicted protein structures was evaluated used Ramachandran plots generated via PROCHECK, assessing the stereochemical parameters to verify proper folding and structural integrity [38]. Model reliability was further determined by calculated DOPE (Discrete Optimized Protein Energy) scores using Modeller, with models exhibit the highest proportion of residues in favored regions and the lowest DOPE scores selected for downstream analyses [35]. The selected models were visualized in UCSF Chimera to assess overall fold correctness and active site architecture. To investigate structural similarity with known plastic-degrading enzymes, the candidate models were superimposed onto crystal structures of characterized PETases using UCSF Chimera [39]. Structural alignment specifically targeted the catalytic triad and adjacent substrate-binding regions [36, 40]. Among the evaluated models, ALF00495.1 and WOZ56011.1 exhibited high structural concordance with PETase reference proteins, supporting their potential functional relevance.

### 2.5. Protein Domain Analysis

The structural features selection, such as α/β-hydrolase folds, PETase-like domains, esterase motifs, and cutinase-specific signatures, the domain architecture of predicted protein sequences was analyzed using InterProScan[41], Pfam databases [42], AlphaFold, and ProSA-web through identified domain and facilitated the classification of proteins into established hydrolytic enzyme families, thereby supporting the prediction of their enzymatic functions [45]. Additionally, comparative domain organization analysis was performed to evaluate the structural and functional similarity of the novel candidate proteins to previously characterized PET-degrading enzymes.

### 2.6. Molecular Dynamics (MD) Simulation

To evaluate the structural stability and dynamic behavior of the selected enzyme models, molecular dynamics (MD) simulations were conducted using GROMACS (version 24.03) [32, 46]. Each protein model was embedded in a cubic simulation box solvated with TIP3P water molecules and neutralized with appropriate counterions. The system underwent energy minimization, followed by equilibration under NVT and NPT ensembles, and subsequent production MD runs. Simulations were performed under standard thermodynamic conditions of 300 K temperature and 1 atm pressure. Simulation analyses included calculation of Root Mean Square Deviation (RMSD), Root Mean Square Fluctuation (RMSF), hydrogen bond, and radius of gyration (Rg) to assess conformational stability and flexibility throughout the simulation trajectory of 100ns[47].

## 3. Results

### 3.1 Target sequences identification

Given that the catalytic triad and domain architecture are critical for maintaining the hydrolytic activity of PET-degrading enzymes[48], we screened a curated dataset of 228 sequences and identified two promising candidates: ALF00495.1 and WOZ56011.1. Sequence analysis confirmed the presence of conserved α/β-hydrolase domains and catalytic residues essential for ester bond cleavage, consistent with the structural framework of known PETases (PlasticDB ID: 00188). The three-dimensional (3D) structures of ALF00495 and WOZ56011 were predicted using AlphaFold, a deep learning-based protein structure prediction tool known for high accuracy [49].

These 3D models revealed conserved PETase-like folds, further validated through structural superimposition with the PETase reference structure. As shown, ALF00495 (**Fig. 1A**) demonstrated closer backbone alignment and superior structural overlap with the template, indicating stronger conservation of the PETase fold. In contrast, WOZ56011 (**Fig. 1B**) displayed comparatively larger deviations, although it retained the overall α/β-hydrolase topology. These results suggest that while both enzymes are structurally consistent with PET-hydrolyzing activity, ALF00495 exhibits greater homology to the reference PETase and represents a more promising candidate for further structural validation and functional studies.

**Figure. 1.**
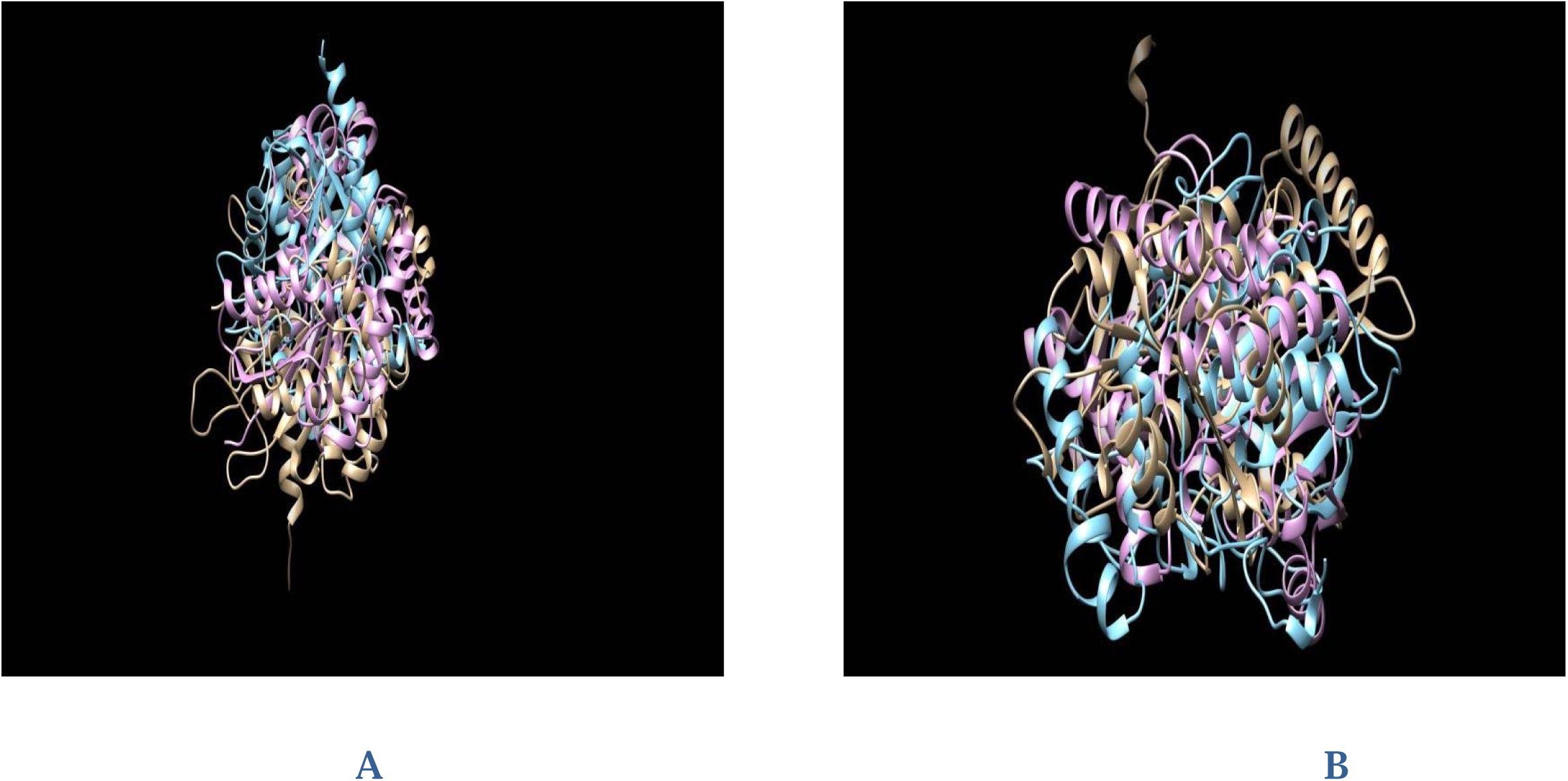
Superimposition structures of model of ALF00495 (A) and WOZ56011 (B). The PETase reference structure is shown in light beige, while the initial ALF00495 (A) and WOZ56011 (B) homology model is colored in light blue.

### 3.2 Model proteins identification and Stability Assessment

Evaluating homology models based on atomic interactions and structural compactness provides critical insight into their energetic favourability and functional reliability[50]. In this study, homology models for two candidate enzymes, ALF00495 and WOZ56011, were constructed using the *Modeller* tool, and their quality was quantified by assessing the Discrete Optimized Protein Energy (DOPE) score, a statistical potential that reflects the model’s overall energy profile. The DOPE score serves as an indicator of energetic stability, with lower values corresponding to fewer steric clashes and more favourable atomic packing[51]. Different models suggested, ALF00495 exhibited a lower DOPE score (−29,650.35), suggesting a more stable structural conformation with minimized regions of high potential energy or steric hindrance. In contrast, WOZ56011 demonstrated a higher DOPE score (−28,810.38), indicating comparatively lower energetic favourability. Structural superimposition with the PETase reference further revealed that ALF00495 (**Fig. 1A**) displayed superior backbone alignment and closer structural overlap with the template, reflecting a higher degree of conservation of the PETase (PlasticDB ID: 00188) fold. WOZ56011 (**Fig. 1B**) although retaining the general α/β-hydrolase topology, showed larger deviations from the reference structure, implying potential differences in flexibility or domain organisation. These results suggest that both enzyme models adopt PETase-like structural frameworks consistent with ester bond hydrolysis, yet ALF00495 presents greater structural fidelity and energetic stability. Consequently, ALF00495 emerges as a more robust candidate for downstream validation.

### 3.3 Structural Superimposition and Validation

Sequence alignment, coupled with curated metadata analysis, confirmed the microbial origin of these enzymes, while phylogenetic profiling underscored their evolutionary association with environments characterised by elevated temperatures. These findings align with the enzyme’s prospective utility in industrial biocatalysis, particularly for polymer degradation under harsh conditions [52]. Superimposition of the modelled structures against the PETase (PlasticDB ID: 00188) reference template revealed conserved overall folds, specifically within the α/β-hydrolase core domain, which is essential for substrate binding and catalysis[53]. Both models retained the hallmark architecture of PET-degrading enzymes; however, ALF00495 (Model A) displayed superior structural alignment, showing more cohesive backbone overlap with the template (**Fig. 1A**). In comparison, WOZ56011 (Model B) exhibited more pronounced deviations, although the general fold remained intact (Fig. 1B). The final selected models, chosen based on their lowest DOPE scores, are represented in light magenta within the superimposition figures (Fig. 2). Furthermore, stereochemical quality was further validated using Ramachandran plot analysis. ALF00495 (Model A) demonstrated excellent geometric parameters, with over 90% of residues located in the most favored regions, 8% in additionally allowed regions, and fewer than 2% in disallowed zones (**Fig. 2A**). Conversely, WOZ56011 (Model B) exhibited a slightly lower stereochemical profile, with approximately 82% of residues in favored regions, 12% in allowed regions, and 6% in disallowed regions (**Fig. 2B**). Despite this modest reduction in quality, both models satisfied the structural benchmarks commonly accepted for homology-modeled proteins. The favorable residue distribution in ALF00495 (Model A), in particular, supports its structural plausibility and suitability for further functional and dynamic characterization.

**Figure. 2.**
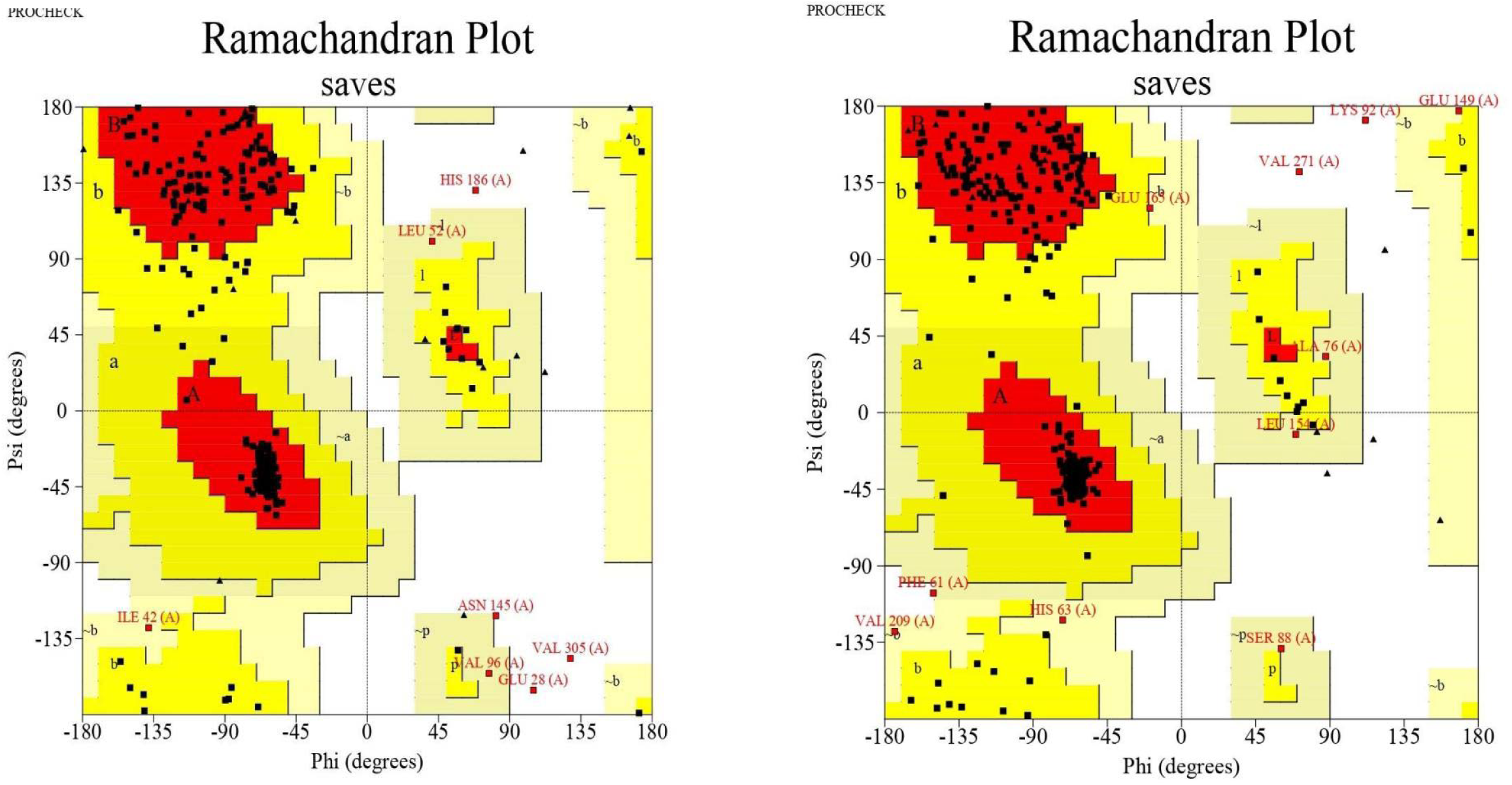
Ramachandran plot validation of stereochemical geometry. Model A (ALF00495) exhibits 90% residues in the most favored regions,whereas Model B (WOZ56011) displays a higher proportion of residues in allowed and disallowed regions, indicating reduced stereochemical quality.

### 3.4 Structure Quality Assessment Based on Z-score Analysis

To determine the structural validity and global fold quality of the modeled PETase-like enzymes ALF00495 and WOZ56011, we conducted comprehensive assessments using the ProSA-web server. The Z-score analysis benchmarks the energy profile of modeled structures against a database of high-resolution experimental structures derived from X-ray crystallography and NMR spectroscopy. ALF00495 exhibited a Z-score of −1.68(**Fig. 3A**), while WOZ56011 scored −1.20(**Fig. 3B**), both values falling within the acceptable range typically observed for native proteins of comparable size and structural class. These scores indicate that the generated models maintain an energetically favorable and physically plausible conformation, can be harnessed for downstream structural and functional studies. In addition to Z-score analysis, we conducted a detailed structural quality evaluation, including residue-wise energy plots and model compactness metrics. On the other hand, the energy distribution across the protein chains was largely favorable for both models, with (**Fig. 4A**) ALF00495 demonstrating more consistent low-energy regions throughout the sequence, particularly in the mid and C-terminal domains. Minimal instances of highly positive energy outliers were observed, further supporting the stability of the predicted folds. Together, these results confirm the structural soundness of the homology models. ALF00495, in particular, demonstrates greater overall energetic stability and structural uniformity. These characteristics not only validate its stereochemical plausibility but also enhance its candidacy for subsequent molecular docking, substrate-binding studies, and catalytic activity simulations.

**Figure. 3.**
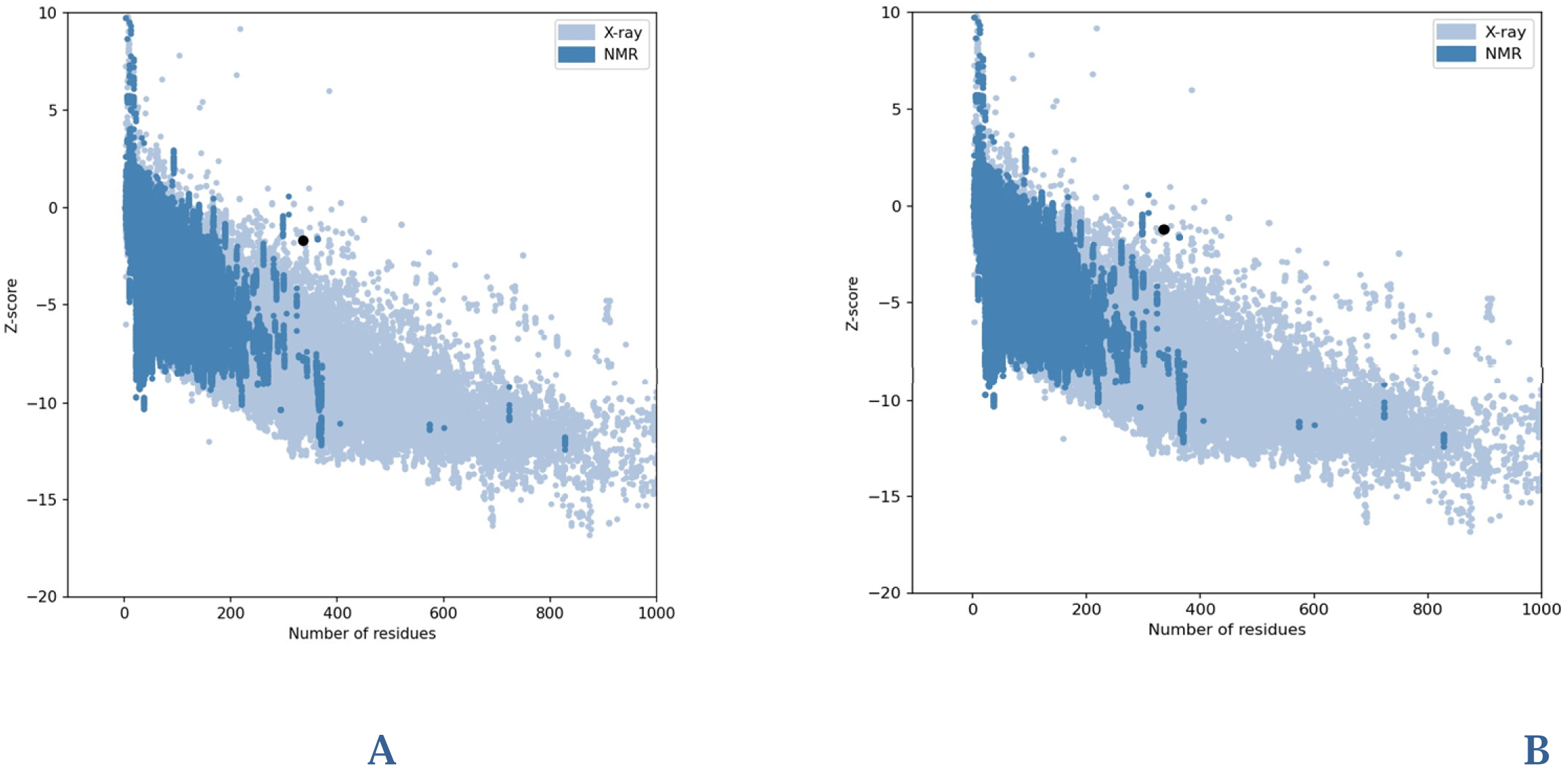
Z-score analysis from ProSA-web server. Both models fall within the acceptable range for native structures; however, Model A demonstrates a more favorable (more negative) Z-score, suggesting higher overall model quality.

**Figure. 4.**
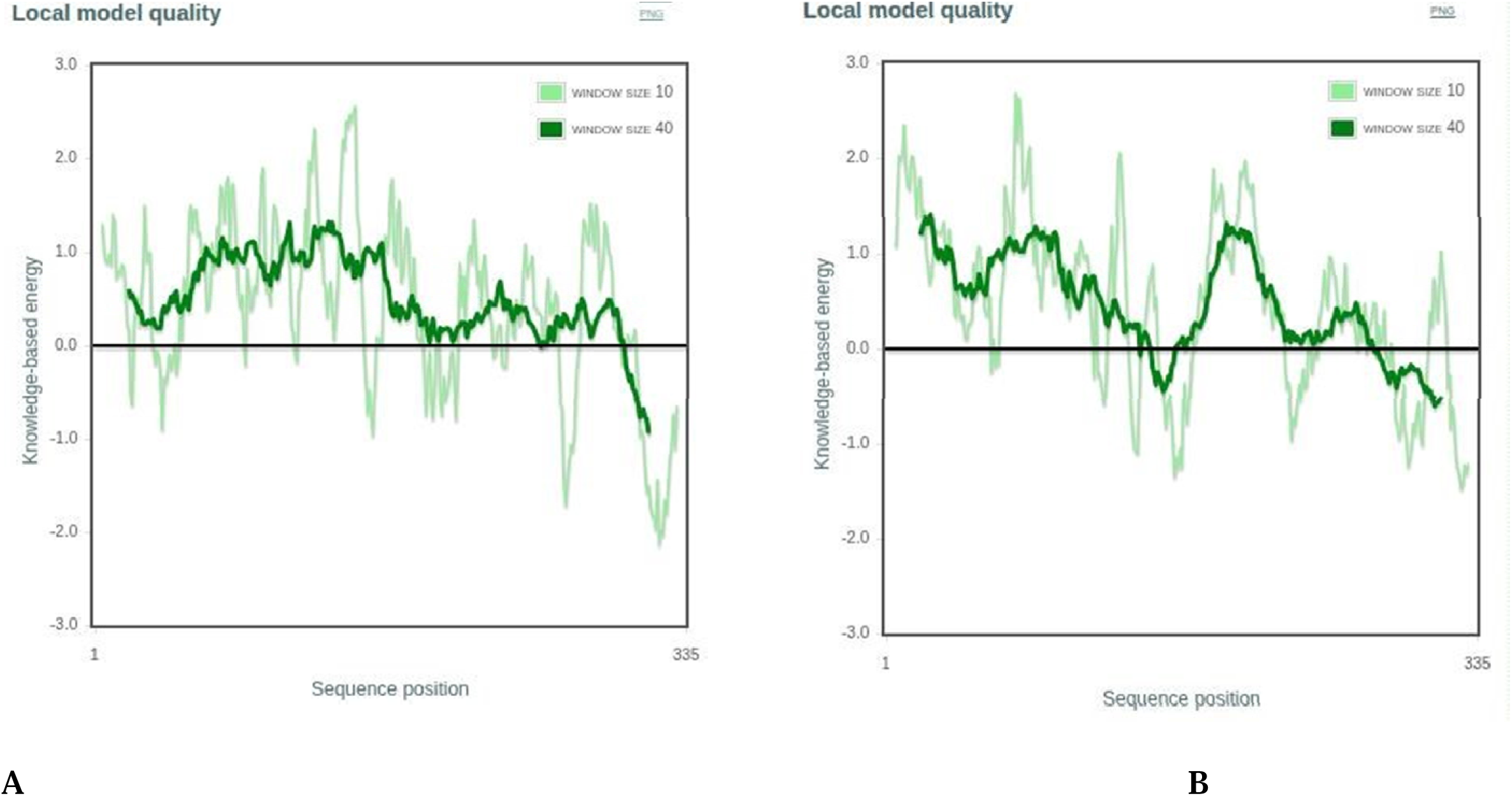
Residue-wise energy profile of modeled structures. Model A (ALF00495) exhibits a smoother energy distribution and fewer high-energy regions compared to Model B (WOZ56011), indicating improved local structural stability.

### 3.5 Molecular Dynamics (MD) Simulation Analysis

Upon exposure to a 100 ns molecular dynamics simulation under aqueous conditions, both ALF00495 (Model A) and WOZ56011 (Model B) showed different dynamic patterns, as evaluated by root-mean-square deviation (RMSD) trajectories.

The structural evolution of ALF00495 (T4) was particularly notable, with its RMSD curve stabilising shortly after the 20 ns mark and maintaining a relatively narrow fluctuation range centered around 0.6 nm for the remainder of the simulation (**Fig. 5, top panel)**. This early convergence indicates a rapid equilibration and reflects the inherent structural stability of the model under physiological conditions. Conversely, the backbone RMSD of WOZ56011 (T13) displayed more irregular fluctuations, particularly during the initial 40 ns, suggesting a delayed accommodation to the simulated environment and increased structural flexibility. Although stabilisation was eventually observed, the amplitude and variability of the RMSD were consistently higher than those of ALF00495, implying comparatively reduced conformational robustness. To further assess the reproducibility of ALF00495’s dynamic behavior, three independent replicate simulations were performed. As shown in the lower panel of (**Fig. 5**), all replicate trajectories (T4_R1 and T4_R2) exhibited a tightly clustered RMSD range near 0.6 nm. This high degree of convergence across simulations substantiates the stability of ALF00495’s fold and supports the notion of its structural reliability during extended simulations. Together, these results reinforce the dynamic resilience of ALF00495 and confirm its suitability for downstream analyses. The sustained low RMSD values and early plateau phase highlight ALF00495’s conformational integrity, while the higher variability observed in WOZ56011 suggests a more flexible, less tightly packed structure that may affect its functional reliability.

**Figure. 5.**
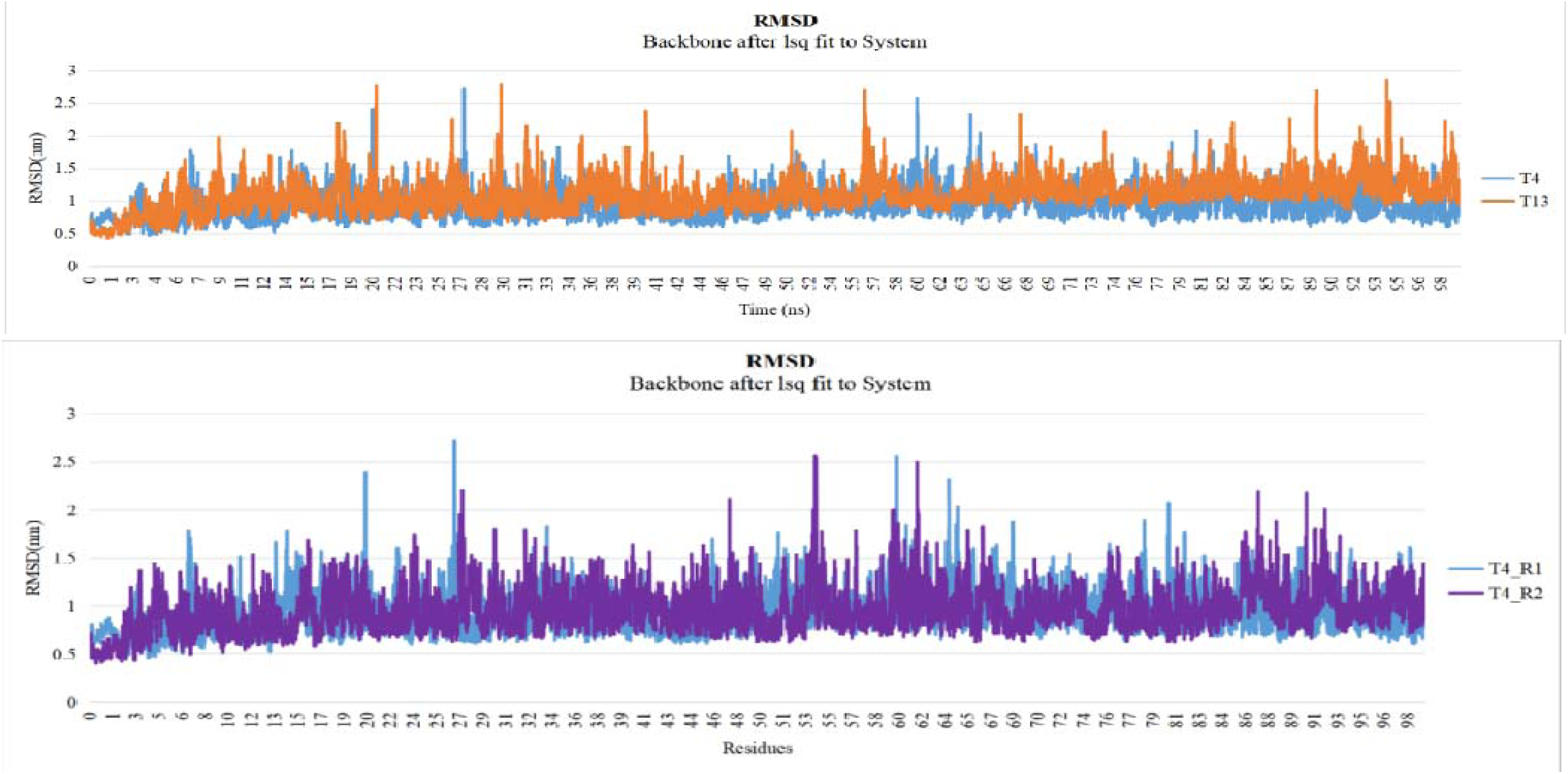
RMSD profile during 100 ns MD simulations. ALF00495 (T4) maintains stable RMSD values (∼0.6 nm) after 40 ns, while WOZ56011 (T13) shows greater fluctuations. Replicate runs (T4_R1–T4_R2) confirm the stability of ALF00495 under physiologi al conditions.

### 3.6 Root Mean Square Fluctuation and Protein Flexibility

To further explore the dynamic behavior and residue-level flexibility of the modeled enzymes, RMSF analysis was conducted for both ALF00495 (Model A) and WOZ56011 (Model B) over a 100 ns molecular dynamics simulation. RMSF values reflect the average displacement of each residue relative to its mean position, thereby highlighting flexible and rigid regions within the protein structure. As presented in (**Fig. 6**), both models exhibited characteristic RMSF patterns, with terminal residues showing higher fluctuation amplitudes, a typical trait in globular proteins due to their increased exposure to solvent. The overall RMSF profile remained within a modest range, spanning from approximately 0.10 to 0.35 nm, indicating stable yet flexible domains. In ALF00495, elevated fluctuations were observed in a few loop regions and surface-exposed residues, particularly between residues 60–85 and 190–210 (**Fig. 6A**). This behaviour suggests potential conformational breathing motions that may facilitate substrate accommodation or allosteric regulation. WOZ56011, on the other hand, demonstrated slightly reduced fluctuations across the majority of its sequence, implying a more rigid structural conformation under simulated conditions (**Fig. 6B**). Notably, in both models, the catalytic core residues critical to PETase activity retained minimal RMSF values (<0.15 nm), signifying stable maintenance of the active site geometry throughout the simulation period. These observations align with the hypothesis that ALF00495, despite its slightly increased local flexibility, preserves essential rigidity within functional regions. The combination of moderate global flexibility and a highly stable catalytic core implies that ALF00495 may exhibit favorable dynamic adaptability, a trait often associated with enhanced substrate binding efficiency and catalytic performance. WOZ56011’s rigidity, while indicative of structural compactness, may limit dynamic responses necessary for flexible substrate interactions. Therefore, RMSF analysis reinforces the dynamic competence of ALF00495 and supports its suitability for downstream functional assays

**Figure. 6.**
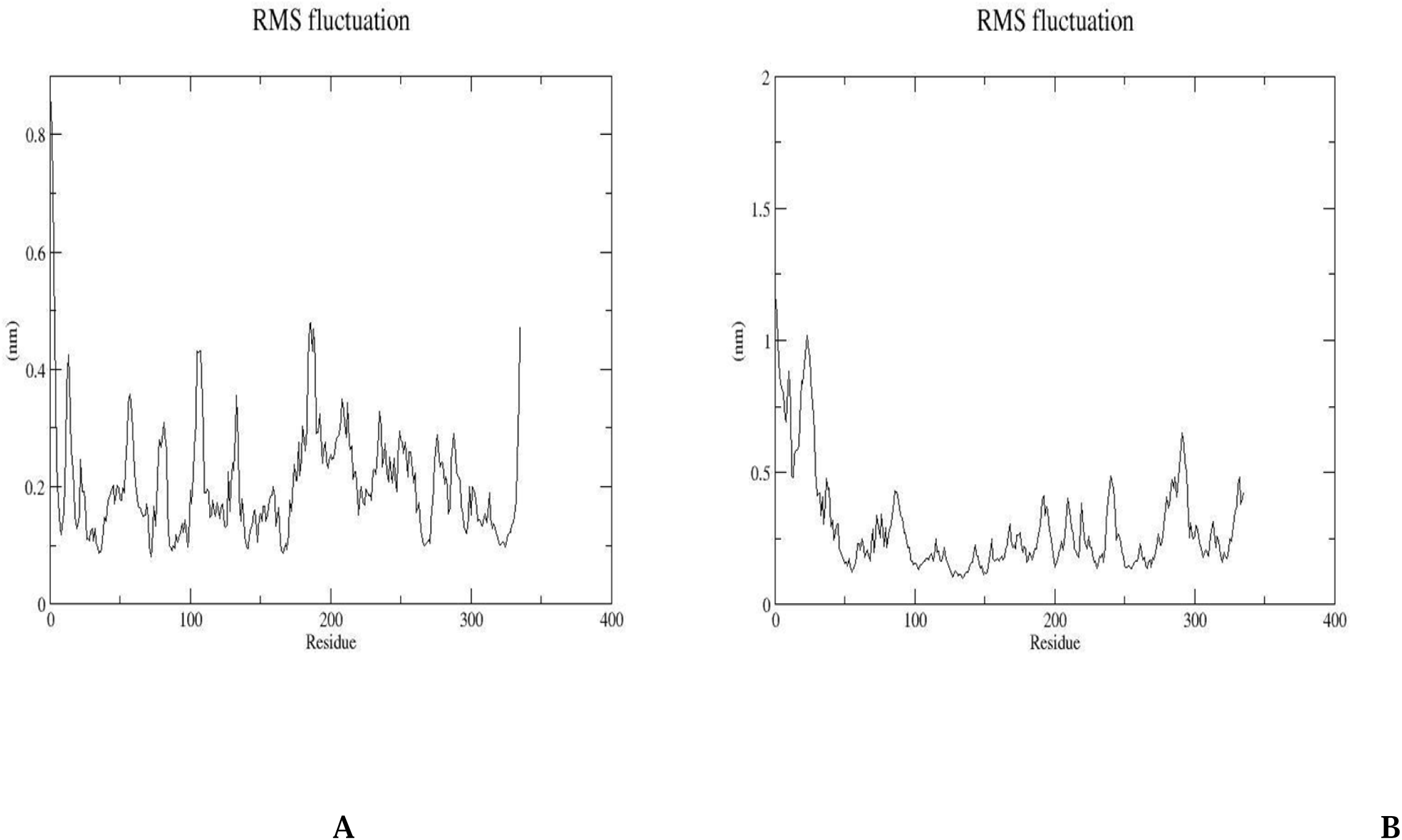
RMSF analysis of atomic-level fluctuations. ALF00495 (Model A) exhibits reduced flexibility across backbone residues, articularly within the α/β-hydrolase domain, while WOZ56011 (Model B) displays increased mobility in terminal and loop regions.

### 3.7 Hydrogen Bond Analysis

Hydrogen bonding is a key determinant of protein structural integrity and functional resilience, especially under physiologically dynamic conditions[54]. To evaluate the intramolecular stability of the modeled PETase-like enzymes, ALF00495 (Model A) and WOZ56011 (Model B), hydrogen bond formation was monitored over a 100 ns molecular dynamics (MD) simulation. As illustrated in (**Fig. 7**), both models demonstrated time-dependent fluctuations in hydrogen bond numbers, indicative of conformational flexibility in an aqueous environment. ALF00495 consistently formed a greater number of hydrogen bonds, averaging between 800 and 900, with initial peaks exceeding 900 during the early simulation phase (0–40 ns) (**Fig. 7A**). This suggests an early structural rearrangement followed by a stable bonding pattern. In contrast, WOZ56011 (**Fig. 7B**) showed slightly lower hydrogen bond counts, fluctuating mostly between 750 and 850, with fewer pronounced peaks, indicating relatively lower structural compactness. Beyond 40 ns, both systems exhibited stabilized hydrogen bonding trends, yet ALF00495 maintained a consistently higher hydrogen bond density throughout the simulation. This persistent intramolecular bonding points to enhanced internal cohesion and superior thermodynamic stability for ALF00495. Collectively, these results reinforce the notion that ALF00495 exhibits stronger internal interactions than WOZ56011, potentially translating into improved performance in substrate binding and catalytic function.

**Figure. 7.**
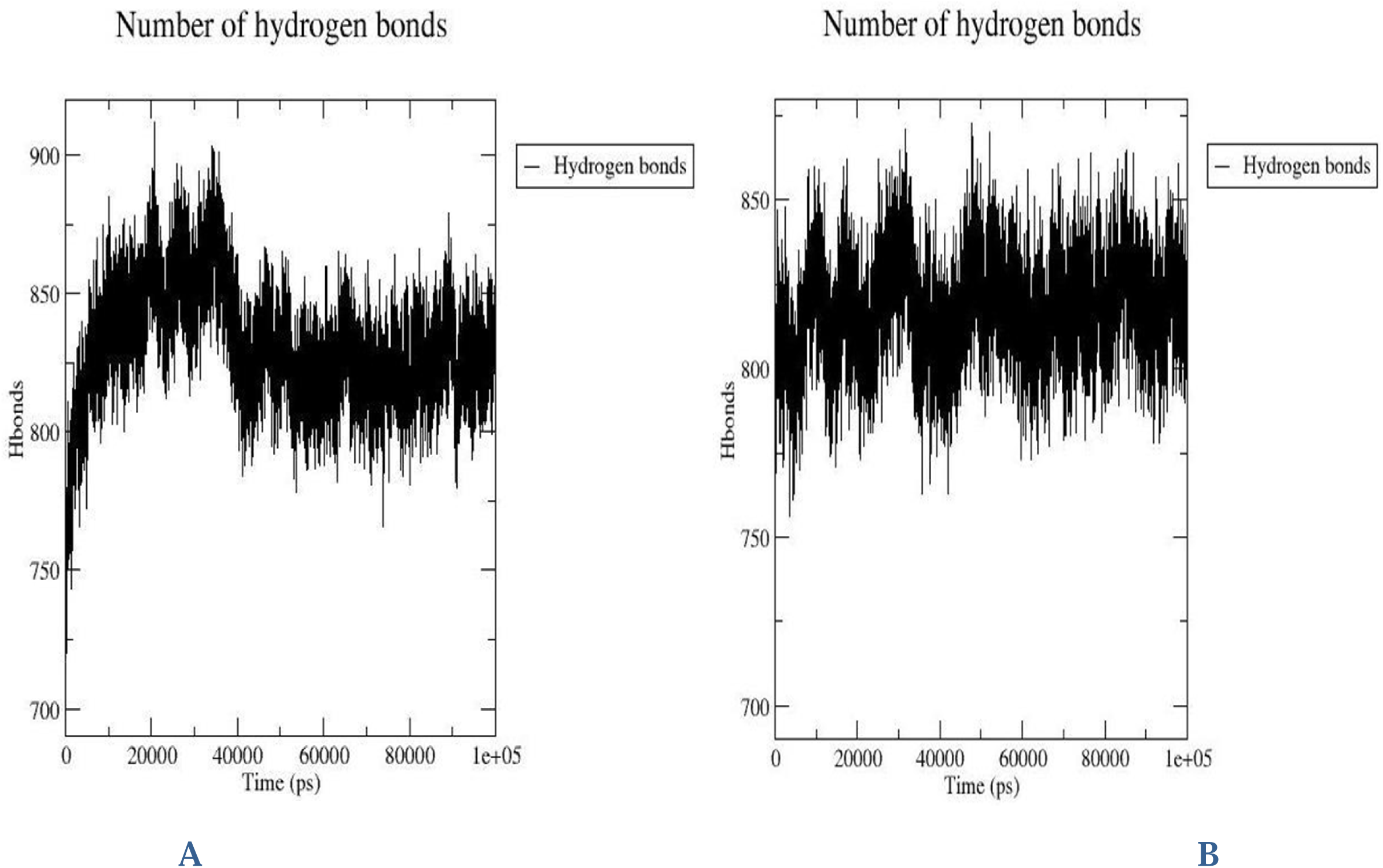
Hydrogen bond formation during MD simulations.

### 3.8 Analysis of Solvent Accessible Surface Area (SASA)

To understand the dynamic behavior of surface exposure in the modeled enzymes, the solvent accessible surface area (SASA) was calculated over the 100 ns molecular dynamics trajectory. The SASA values were analyzed to correlate changes in surface area with conformational flexibility and potential functional accessibility[55]. For ALF00495, the SASA gradually increased during the initial 40 ns, reaching a steady range between 210 and 220 nm² (**Fig. 8A**). This trend indicates an expansion of the molecular surface, potentially linked to dynamic loop rearrangements or unfolding of peripheral domains. The surface area remained stable in the latter half of the simulation, suggesting structural equilibrium in a moderately solvent-exposed conformation. In contrast, WOZ56011 exhibited a biphasic SASA profile. An initial increase was observed, followed by a gradual decrease in surface exposure after 50 ns, eventually stabilizing below 200 nm² (**Fig. 8B**). This behaviour implies a compaction event, where flexible regions possibly folded inward, resulting in a more shielded and compact structure. These distinct SASA profiles reflect differing structural responses under identical simulation conditions. The increasing SASA of ALF00495 may allow greater interaction potential with solvent or substrates, while the reduction in WOZ56011 suggests a conformation favouring structural rigidity. These findings support the utility of SASA as a reliable metric for evaluating protein surface dynamics and highlight ALF00495’s potential as a more interaction-competent candidate.

**Figure. 8.**
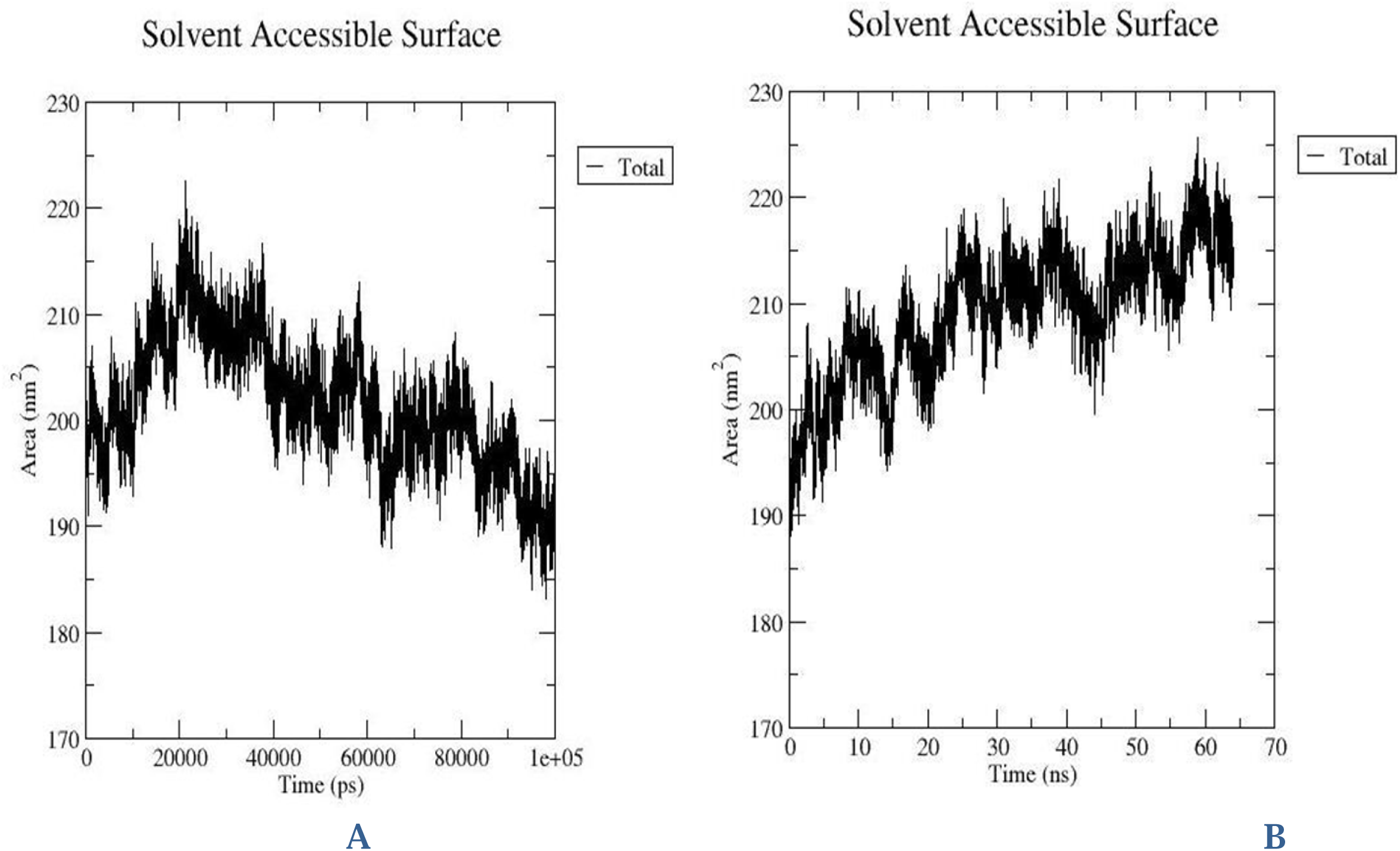
Solvent accessible surface area (SASA) over 100 ns simulation. ALF00495( Model A) displays a lower and more stable SASA profile, reflecting a compact fold, while WOZ56011( Model B) exhibits relatively higher solvent exposure and variability.

### 3.9 Radius of Gyration and Structural Compactness

To assess the global folding behavior and compactness of the modeled PETase-like enzymes, the radius of gyration (Rg) was computed for both ALF00495 (Model A) and WOZ56011 (Model B) across the 100 ns molecular dynamics simulation. Rg is a key indicator of a protein’s structural compactness, defined as the mass-weighted root-mean-square distance of the atoms from their common center of mass. As illustrated in (**Fig. 9A**), ALF00495 exhibited a consistent Rg trajectory ranging between 2.0 and 2.1 nm throughout the entire simulation, indicating stable structural compactness. The minimal fluctuation in Rg values reflects a tightly packed conformation and a low degree of structural expansion, suggesting that the model maintained its folded state with high spatial fidelity under simulated physiological conditions. In contrast, (**Fig. 9B**) WOZ56011 displayed a slightly broader range of Rg values, initially increasing during the first 30 ns of simulation before stabilizing between 2.15 and 2.25 nm. This pattern suggests an initial phase of structural rearrangement followed by equilibration. The comparatively larger Rg values of WOZ56011 imply a more relaxed and extended conformation, possibly linked to the higher RMSD and reduced stereochemical quality observed in prior analyses. The consistently lower Rg values observed for ALF00495 reinforce its compact and stable tertiary architecture, a characteristic often associated with thermophilic proteins. Such compactness enhances thermal stability by minimising solvent exposure and internal cavity formation, which are crucial factors in maintaining functional integrity under fluctuating environmental conditions. Overall, the Rg analysis further substantiates ALF00495’s superior structural integrity and compactness. These findings, in conjunction with RMSD and RMSF results, position ALF00495 as a dynamically stable and structurally favourable candidate for further PETase-related functional investigations.

**Figure. 9.**
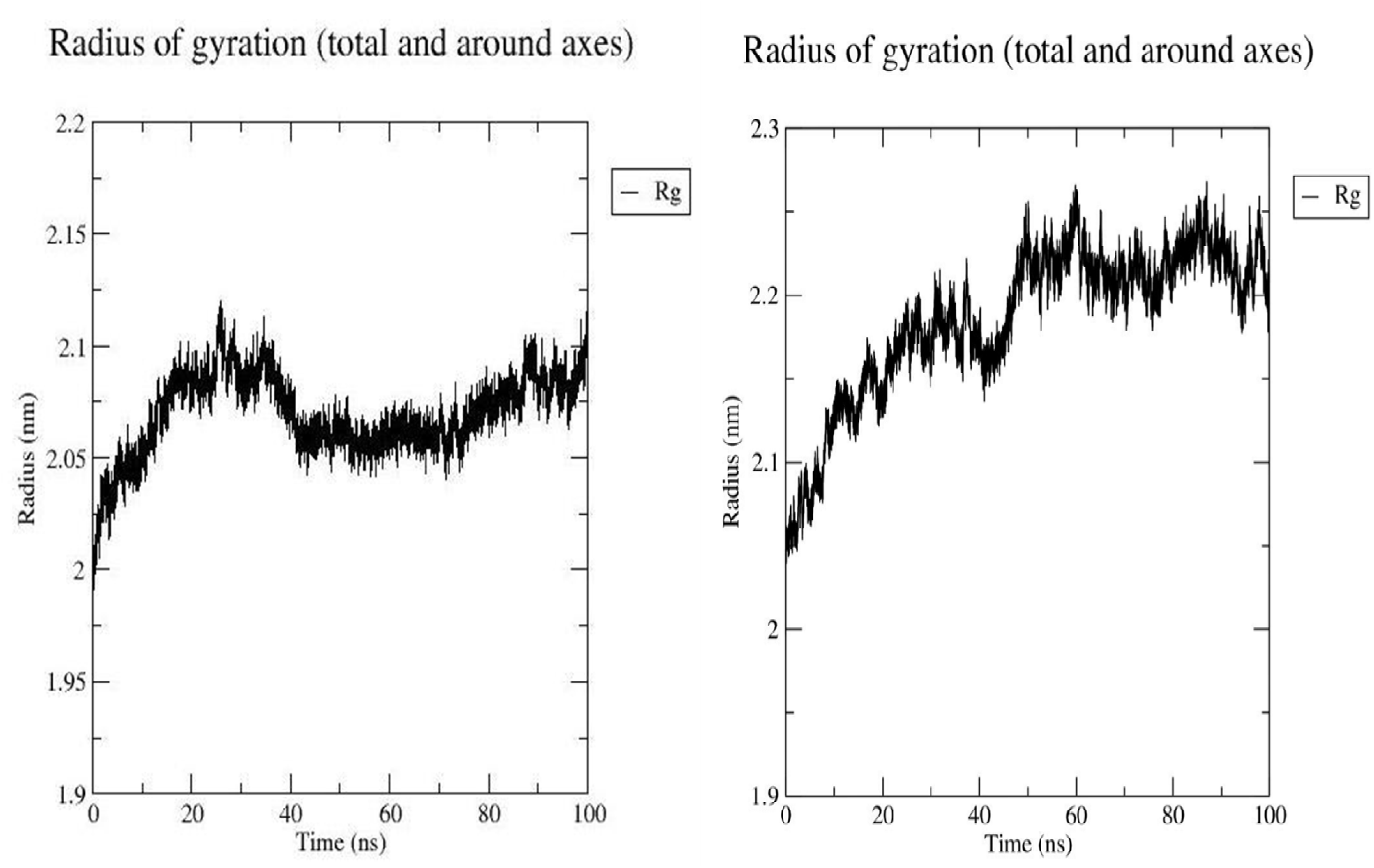
Radius of gyration (Rg) during simulation. ALF00495 (Model A) maintains lower Rg values, suggesting enhanced compactness and conformational stability. WOZ56011(Model B) shows slight expansion over time.

## 4. Discussion

The environmental burden of polyethylene terephthalate (PET) accumulation has accelerated efforts to identify and optimize enzymes capable of efficient PET degradation. Among these, *Ideonella sakaiensis* PETase has received significant attention for its ability to depolymerize PET under mild conditions [56]. However, its limited thermal stability and modest catalytic rates constrain industrial translation [57]. This has prompted the search for PETase variants or novel homologs with enhanced robustness and activity. Thermophilic actinobacteria, including *Thermobifida* species, represent attractive reservoirs, as enzymes from thermophiles often display superior stability and are naturally adapted to harsh environments that may favor efficient PET hydrolysis[58].

Son et al. (2018) reported that PETase-like enzymes adopt an α/β-hydrolase fold with the canonical Ser–His–Asp catalytic triad, supporting our classification of ALF00495 and WOZ56011 as PETase-like enzymes. The structural predictions using AlphaFold further revealed that ALF00495 closely aligned with the canonical PETase(PlasticDB id: 00188) backbone, while WOZ56011 retained the overall fold but exhibited larger deviations[49]. This suggested that ALF00495 may preserve functional homology more faithfully, whereas WOZ56011 could represent a structurally divergent variant with altered dynamics [49]. The quality of the modeled structures was evaluated using ProSA-web, which confirmed energetically favorable conformations for both enzymes (Z-scores 1.68 for ALF00495 and −1.20 for WOZ56011) within the range of native proteins. Notably, ALF00495 demonstrated more uniform residue-wise energy distribution and fewer high-energy outliers, reinforcing its structural plausibility. Together, these results validated both models while highlighting ALF00495 as the more robust candidate [59].

Molecular dynamics (MD) simulations provided deeper insights into the conformational resilience of these enzymes under aqueous conditions. Root-mean-square deviation (RMSD) trajectories revealed early equilibration and sustained stability for ALF00495, in contrast to the delayed stabilization and higher fluctuations of WOZ56011. Replicate simulations of ALF00495 confirmed reproducibility, further underscoring its dynamic stability. These findings are consistent with prior reports linking conformational robustness to higher catalytic efficiency in PETases[60]. Root-mean-square fluctuation (RMSF) analysis offered residue-level perspectives on flexibility[46, 61]. Teodoro et al. (2003) reported that terminal regions of globular proteins typically exhibit higher flexibility, whereas Bartlett et al. (2002) demonstrated that catalytic residues remain structurally rigid to preserve active-site geometry, consistent with our observations in both models. Importantly, ALF00495 displayed localized flexibility in surface loops suggesting adaptive motions that may facilitate substrate access or allosteric adjustments[62]. WOZ56011, by contrast, showed reduced overall fluctuations, reflecting structural rigidity[63]. While rigidity may enhance fold stability, excessive restriction could limit dynamic adaptability required for efficient PET binding and turnover [64].

Hydrogen bonding analysis further differentiated the two models[65]. ALF00495 consistently formed more intramolecular hydrogen bonds, particularly during the early simulation phase, where an initial surge was followed by stabilization into a dense network. This behavior suggests enhanced thermodynamic stability and compact folding, attributes strongly correlated with enzyme resilience under dynamic conditions[66, 67]. In comparison, WOZ56011 maintained fewer hydrogen bonds, indicative of reduced compactness and weaker internal cohesion[65]. Solvent accessible surface area (SASA) analysis revealed contrasting dynamic responses. ALF00495 showed a gradual increase in surface exposure, stabilizing at higher values that may support greater substrate accessibility. WOZ56011, however, underwent a compaction event, reducing SASA and suggesting a more shielded structure.While this may contribute to stability, it likely restricts functional adaptability. Together, these results suggest that ALF00495 balances compactness with moderate flexibility, a combination often favorable for catalysis [68, 69]. The radius of gyration (Rg) profiles reinforced these observations. ALF00495 maintained consistently low Rg values (∼2.0–2.1 nm), reflecting a compact, thermophilic-like fold with minimal structural expansion[70]. In contrast, WOZ56011 exhibited higher and more variable Rg values, consistent with a looser conformation and reduced structural integrity[71]. Compactness is a hallmark of stable enzymes adapted to elevated temperatures[72, 73], and the lower Rg values of ALF00495 align with its superior stereochemical quality and dynamic resilience observed across analyses.

Integrating these findings, ALF00495 emerges as the stronger PETase-like candidate, combining favorable stereochemical quality, energetic stability, compact folding, and adaptive flexibility with a rigid catalytic core[30, 74, 75]. These attributes suggest not only robustness but also functional competence, positioning ALF00495 as a promising scaffold for downstream applications, including molecular docking, substrate-binding assays, and protein engineering aimed at enhancing PET degradation[15, 30, 75]. WOZ56011, while less stable, retains the α/β-hydrolase topology and may provide complementary insights into structural variation and its impact on catalytic efficiency [15, 30]. Overall, this work expands the repertoire of PETase-like enzymes by identifying and characterizing two thermophilic candidates. The comprehensive computational evaluation highlights ALF00495 as a particularly promising enzyme for further experimental validation and potential engineering toward scalable, eco-friendly PET recycling strategies.

### Conclusion

*Thermobifida cellulosilytica* and *Thermobifida halotolerans,* thermophilic actinobacteria, are highlighted in this study as potential sources of novel PETase-like enzymes for the degradation of plastic at high temperatures. Two potential enzymes (ALF00495 and WOZ56011) were found and thoroughly verified using an integrated in silico process that combined sequence homology, structural modeling, and molecular dynamics simulations. ALF00495 is a particularly good candidate for further experimental characterization because of its excellent stereochemical quality, improved dynamic stability, and structural compactness.

Their thermostable structural characteristics meet the needs for industrial-scale recycling at high temperatures, and the retention of the traditional Ser–His–Asp catalytic triad and α/β-hydrolase fold in both models validates their possible hydrolytic activity against PET substrates. These results highlight the unexplored enzymatic diversity of thermophilic microbes and confirm the effectiveness of computational methods as an expedient means of enzyme engineering and discovery. Ultimately, this work establishes a strong basis for creating eco-friendly, effective enzymatic recycling systems and hybrid degrading techniques to tackle the expanding global plastic waste problem.

## Statements & Declarations

### Author Contributions

Satyabharati Giri (S.G.) conceived the study and supervised/project administration. Amrendra Kumar (A.K.), Arbaz Attar (A.A.) and Bhagyashree Patil (B.P.) developed the methodology. B.P. and A.K. performed software development and formal analysis. B.P. and A.K. carried out the investigation and curated the data. B.P. prepared the visualizations. B.P., A.K., and S.G. drafted the original manuscript. S.G. reviewed and edited the manuscript.

## Acknowledgements

We thank the SMCS-Psi Data Analytics team for valuable discussions and technical support. We also acknowledge PyMOL.We also used MODELLER for comparative modeling, GROMACS 24.03 for molecular dynamics, AutoDock Vina for docking, RDKit for cheminformatics.

## Data Availability

All data and materials supporting the findings of this study (e.g.,curated PlasticDB sequence set, alignments, homology models, MD input parameters, and analysis scripts) are available from the corresponding author on reasonable request.

## Code Availability

Custom scripts and analysis notebooks are available from the corresponding author on reasonable request.

## Ethics approval / Consent to participate / Consent to publish

Not applicable. This research did not involve human participants, animals, or identifiable personal data.

